# Fenestrae-associated protein Plvap regulates the rate of blood-borne proteins passage into the hypophysis

**DOI:** 10.1101/571299

**Authors:** Ludmila Gordon, Janna Blechman, Eyal Shimoni, Dvir Gur, Bela Anand-Apte, Gil Levkowitz

## Abstract

To maintain body homeostasis, endocrine systems must detect and integrate a multitude of blood-borne peripheral signals. This is mediated by specialized permeable pores in the endothelial membrane, dubbed fenestrae. Plasmalemma vesicles-associated protein (Plvap) is located in the fenestral diaphragm and is thought play a role in the selective passage of proteins through the fenestrae. However, this suggested function has yet to be demonstrated directly. Here, we studied the development of fenestrated capillaries in a major neuroendocrine interface between the blood and brain, namely the hypophysis. Using a transgenic permeability biosensor to visualize the vascular excretion of a genetically tagged plasma protein (DBP-EGFP), we show that the developmental acquisition of vascular permeability is associated with differential expression of zebrafish *plvap* orthologs in the hypophysis versus brain. Ultrastructural analysis of the hypophyseal vasculature revealed that *plvapb* mutants display deficiencies in fenestral and stomatal diaphragms as well as increased density of fenestrae, but not of caveolae. Measurements of DBP-EGFP dynamics in live *plvapb* mutant larvae provided a direct proof that Plvap limits the rate of blood-borne protein passage through fenestrated endothelia. Overall, we present the regulatory role of Plvap in the development of blood-borne protein detection machinery in a major neuroendocrine interface between the brain and the general circulation.

## Introduction

Endocrine organs regulate homeostasis by interfacing with blood vessels to enable the bidirectional passage of peptide hormones and blood-borne molecules to and from the general circulation. The capillaries of all endocrine organs contain fenestrae, which are circular membranal openings that cut through the cell body of endothelial cells. Fenestrae play a principal role in endocrine regulation of homeostasis by allowing higher permeability to small and medium-sized molecules. Fenestrated capillaries are usually found in organs with rapid solute exchange such as endocrine glands, gastrointestinal tract mucosa, and kidney peritubular capillaries [reviewed in (Aird, 2007; Stan, 2007). There are three major types of fenestrae: diaphragmed, non-diaphragmed and sinusoidal.

The diaphragmed fenestrae are pores, typically of 60-70 nm in diameter, that are traversed by a thin diaphragm (Rhodin, 1962). The only known molecular component associated with the fenestrae diaphragm is the type II transmembrane glycoprotein, which is encoded by the vertebrate plasmalemma vesicles-associated protein *(PLVAP)* gene (Stan et al., 1999). Fenestral diaphragms consist of PLVAP homodimers, whose C termini form the central density of the diaphragm [reviewed in (Stan et al., 1999)]. In addition to its presence in the diaphragms of endothelial fenestrae, PLVAP is also localized in stomatal diaphragms of caveolae, as well as in transendothelial channels and endothelial pockets (Hamilton et al., 2019; Milici et al., 1986; Stan, 2005; Stan et al., 1999; Stan et al., 2004). Caveolae, which are flask-shaped invaginations of the plasma membrane of regular shape and size (50-100 nm), are found in endothelial and other cell types and are known to be involved in transcytosis [reviewed in (Filippini et al., 2018; Stan, 2005)].

Several independent studies of *Plvap*-deficient mice have shown that PLVAP protein is required for diaphragm formation; however, it is not required for the formation of fenestral or caveolar pores (Herrnberger et al., 2012; Stan et al., 2012). The impairment of fenestral diaphragms has led to impaired barrier function of fenestrated capillaries, causing a disruption of blood composition, i.e. hypoproteinemia and hypertriglyceridemia. This led to early death of animals due to severe non-inflammatory protein-losing enteropathy, which is caused by uncontrolled plasma proteins leakage (Herrnberger et al., 2012; Stan et al., 2012). It was suggested that PLVAP limits the passage of proteins through the fenestral pore, however, thus far there has been no direct demonstration of this suggested function.

The hypothalamo-hypophyseal system, which consists of the median eminence and neurohypophysis, is a major neuroendocrine interface, which allows the brain to regulate homeostasis by recognizing blood-borne proteins and releasing neurohormones into the general circulation (Anbalagan et al., 2018; Gutnick et al., 2011; Wircer, 2016). These two brain interfaces are also known as circumventricular organs (CVO), which are specialized areas located around the midlines of the brain ventricles (Ganong, 2000; Miyata, 2015). In contrast to CNS vasculature, which contains blood-brain barrier (BBB), CVO’s capillaries are fenestrated and, as such, they facilitate the bidirectional exchange of information between the CNS and the periphery without disrupting the BBB (Ciofi et al., 2009; Ganong, 2000; Miyata, 2015; Schaeffer et al., 2014). The role of PLVAP in CVO permeability, including that of the hypophysis, has never been studied before.

Here, we investigated the necessity of Plvap for the developmental acquisition of permeability in the fenestrated capillaries of the zebrafish hypophysis. We found that the expression of two zebrafish PLVAP orthologs during development coincides with the acquisition of hypophyseal permeability to plasma proteins and the establishment of permeability boundary between the hypophysis and BBB-containing vasculature. We employed live imaging of zebrafish larvae to determine the dynamics by which a genetically-labeled plasma protein passes through hypophyseal endothelia. Finally, we provide the first direct proof that PLVAP regulates the rate of blood-borne protein transfer through fenestrated endothelia into the hypophysis.

## Results

### Hypophyseal vasculature is permeable to blood-borne proteins and lack BBB

To study the regulation of blood-borne protein passage through the hypophyseal blood capillaries, we used the zebrafish as a model system. Unlike in mammals, in which the hypothalamo-hypophyseal system is segregated into median eminence and neurohypophysis, zebrafish hypophyseal vasculature consists of two adjoined neurovascular interfaces, rostral and caudal pars nervosa, that originate from a hypophyseal vascular loop-like structure (Anbalagan et al., 2018; Gutnick et al., 2011; Liu et al., 2013). Hypophyseal capillaries of zebrafish larvae have small diameter and they might be ruptured when using standard micro-angiography due to increased blood volume. Therefore, we employed a non-invasive vascular permeability reporter, Tg(*l-fabp*:DBP-EGFP;*kdrl*:mCherry-caax) (Xie et al., 2010), which we have recently used to demonstrate hypophyseal permeability at 5 days post-fertilization (dpf) (Anbalagan et al., 2018). This double transgenic permeability reporter expresses the vitamin D-binding plasma protein fused to EGFP (DBP-EGFP) and a membrane-tethered mCherry reporter in endothelia, allowing simultaneous visualization of vascular morphology and DBP-EGFP extravasation from the blood into the parenchyma (Fig. 1A).

**Figure 1.**
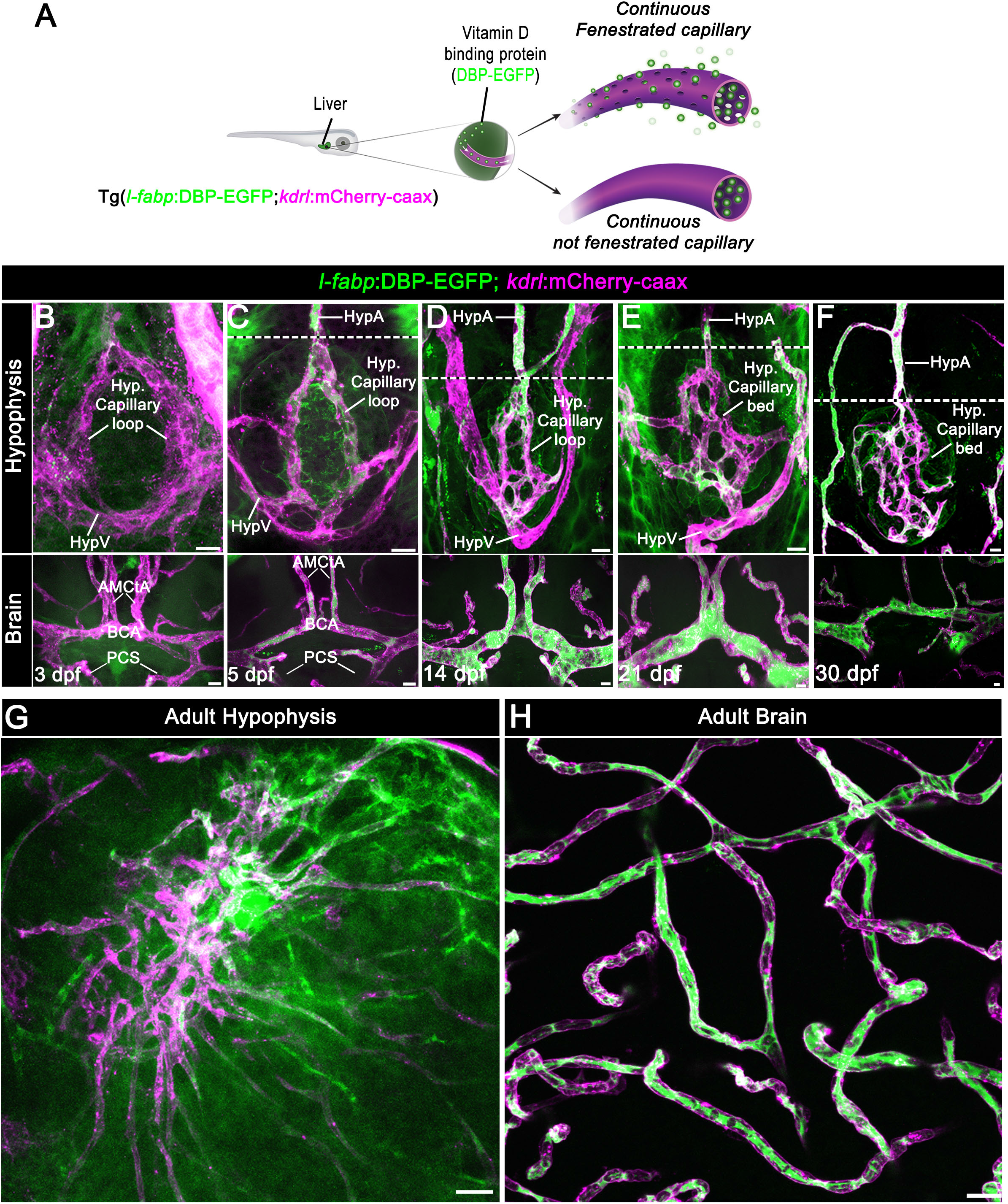
Zebrafish neurohypophyseal vasculature is permeable to blood-borne proteins. **(A)** Scheme describing an endogenous biosensor for real-time monitoring of vascular permeability. Vitamin D-binding protein (DBP) fused to EGFP, expressed in hepatocytes under liver-specific promoter (1-fabp) and secreted into the general circulation, served as permeability biosensor. **(B-F)** Whole-mounts of double transgenic Tg(*l-fabp*:DBP-EGFP;*kdrl*:mCherry-caax) zebrafish at different developmental stages demonstrating excessive extravasation of DBP-EGFP in the pituitary (top) but not in the brain (bottom). Note the functional boundary established between the capillary loop and hypophyseal artery (dotted lines). HypA, hypophyseal artery, AMCtA, Anterior (rostral) mesencephalic central artery, BCA, Basal communicating artery, PCS, Posterior (caudal) communicating segment. Scale bars: 5 μm. **(G,H)** Whole-mounts of double transgenic Tg(*l-fabp*:DBP-EGFP;*kdrl*:mCherry-caax) adult zebrafish hypophysis **(G)** and brain **(H)** vasculature. Scale bars: 20 μm.

First, we visualized the extravasation of DBP-EGFP in the developing hypophyseal vasculature as well as in the adjacent brain vasculature (Fig. 1B-F). As we reported previously (Gutnick et al., 2011), the formation of a simple hypophyseal vascular loop-like structure is initiated at 3 dpf. At this embryonic stage, we detected faint but consistent DBP-EGFP signal inside the hypophyseal loop, as well as outside the hypophyseal artery and brain vasculature (Fig. 1B). At 5 dpf, a more pronounced extravasated DBP-EGFP signal was detected in the whole hypophyseal circumference in and around the hypophyseal capillary loop. At this post-embryonic larval stage, a permeability boundary started to form between the hypophyseal loop and its inferior hypophyseal artery, which extends from the brain and connects to the capillary loop (dotted line in Fig. 1C). Thus, while the hypophyseal loop displayed extensive DBP-EGFP extravasation, the anterior part of the hypophyseal artery and the CNS vasculature retained the DBP-EGFP inside the vascular lumen. This permeability boundary between hypophyseal and brain vasculature became more prominent at juvenile stages (14-30 dpf), during which the hypophyseal capillary loop underwent extensive angiogenic sprouting. Specifically, the simple loop has transformed into an elaborated network of permeable microvessels (Fig. 1D-F) and, thereafter, formed highly complex hypophyseal capillary plexus in the adult fish (Fig. 1G). While the capillary plexus of the hypophysis maintained permeability to blood-borne protein, the brain vessels (Fig. 1H) remained non-permeable. As expected, hypophyseal capillaries of either 5 dpf larvae or adult zebrafish did not express the common BBB-associated tight junction marker, claudin-5, whereas CNS vasculature displayed an extensive anti-claudin-5 staining (Fig. 2). Thus, the functionalization of permeable hypophyseal capillaries inversely correlated with the formation of BBB in CNS vasculature in the developing larvae.

**Figure 2.**
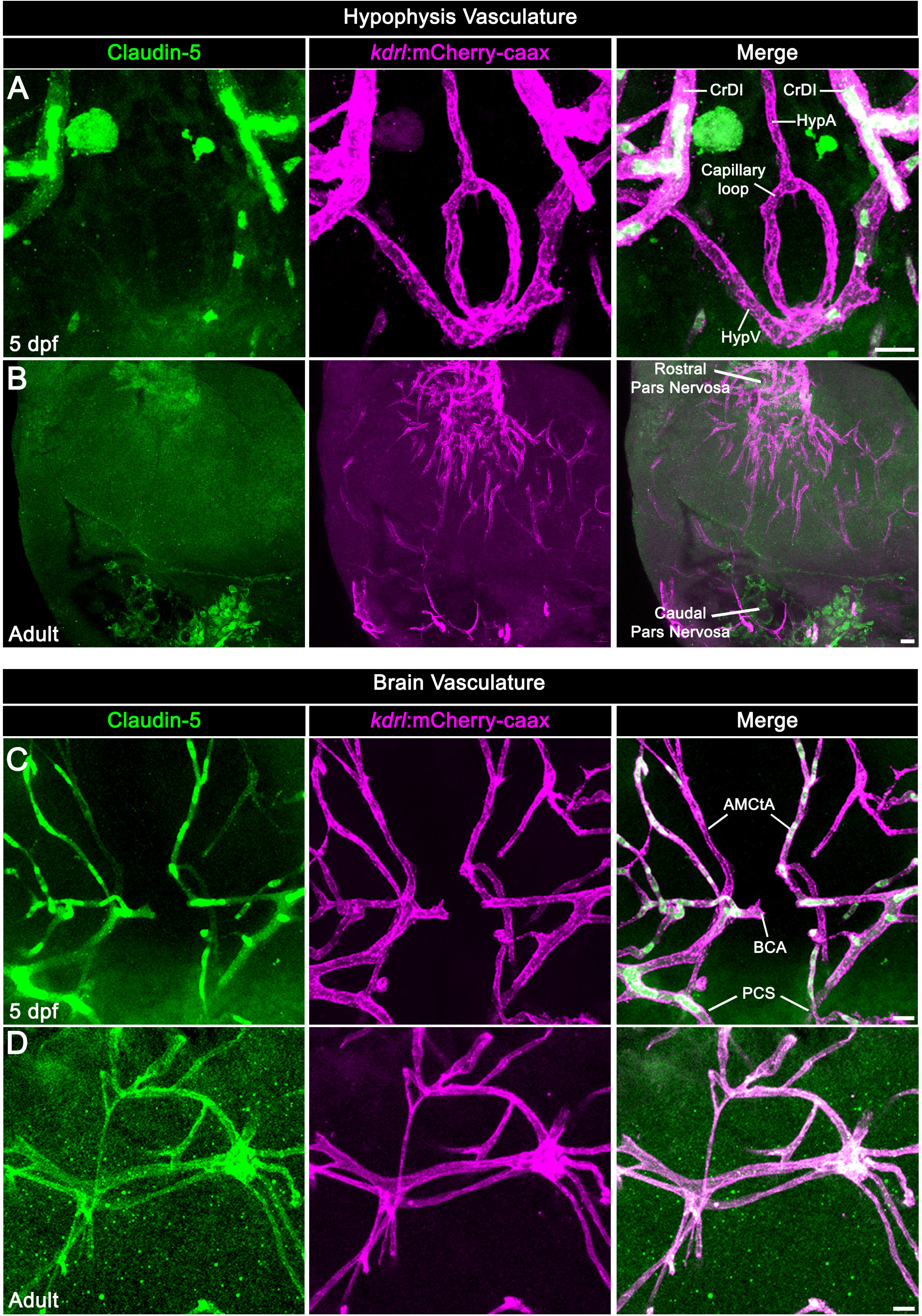
Neurohypophyseal vasculature is devoid of the BBB marker protein claudin-5. (**A-B**) Claudin-5 expression by the hypophyseal vessels was detected by whole-mount immunofluorescent staining using anti-Claudin 5 antibody in 5 dpf larvae (A) and adult (B) Tg(*kdrl*:mCherry-caax) zebrafish. (**C-D**) Claudin-5 expression by the brain vessels was detected by whole-mount immunofluorescent staining using anti-Claudin 5 antibody in 5 dpf larvae (C) and adult (D) Tg(*kdrl*:mCherry-caax) zebrafish. CrDI, Cranial division of the internal carotid artery, HypA, hypophyseal artery, AMCtA, Anterior (rostral) mesencephalic central artery, BCA, Basal communicating artery, PCS, Posterior (caudal) communicating segment. Scale bars: 5 μm (A, C); 20 μm (B, D).

### Zebrafish hypophyseal capillaries contain fenestrae and caveolae

It was reported that vertebrate hypophyseal capillaries contain multiple diaphragmed fenestral openings of about 60-80 nm in size (Farquhar, 1961; Gross et al., 1986; Holmes and Ball, 1974; Stan et al., 2012); however, there are no similar reports in zebrafish. To examine whether the blood vessels in the zebrafish hypophysis are fenestrated, we subjected zebrafish larvae and adults to transmission electron microscopy (TEM) (Fig. 3A,B) and scanning electron microscopy (cryo-SEM). Due to the small size (~50 x 20 μm) of the hypophyseal capillary loop area in the larvae, we used the Tg(*oxtl*:EGFP) (Blechman et al., 2011) as a fluorescent landmark to identify the hypophysis for imaging (Fig. 3A). Our TEM imaging of hypophyseal sections demonstrated that diaphragmed fenestrae exist in the hypophyseal endothelia already in 5 dpf larvae (Fig. 3C). Moreover, both larval and adult hypophyseal capillaries contained diaphragmed fenestrae located along the endothelial wall, which appear as sieve-like structures (Fig. 3D). Finally, using cryo-SEM imaging, we obtained a surface view of the fenestral diaphragms, revealing that these diaphragms are dissected into several openings by fibrils that converge at the center of the pore (Fig. 3G). We conclude that zebrafish hypophyseal endothelium is fenestrated already at early developmental stages and remains fenestrated in adulthood.

**Figure 3.**
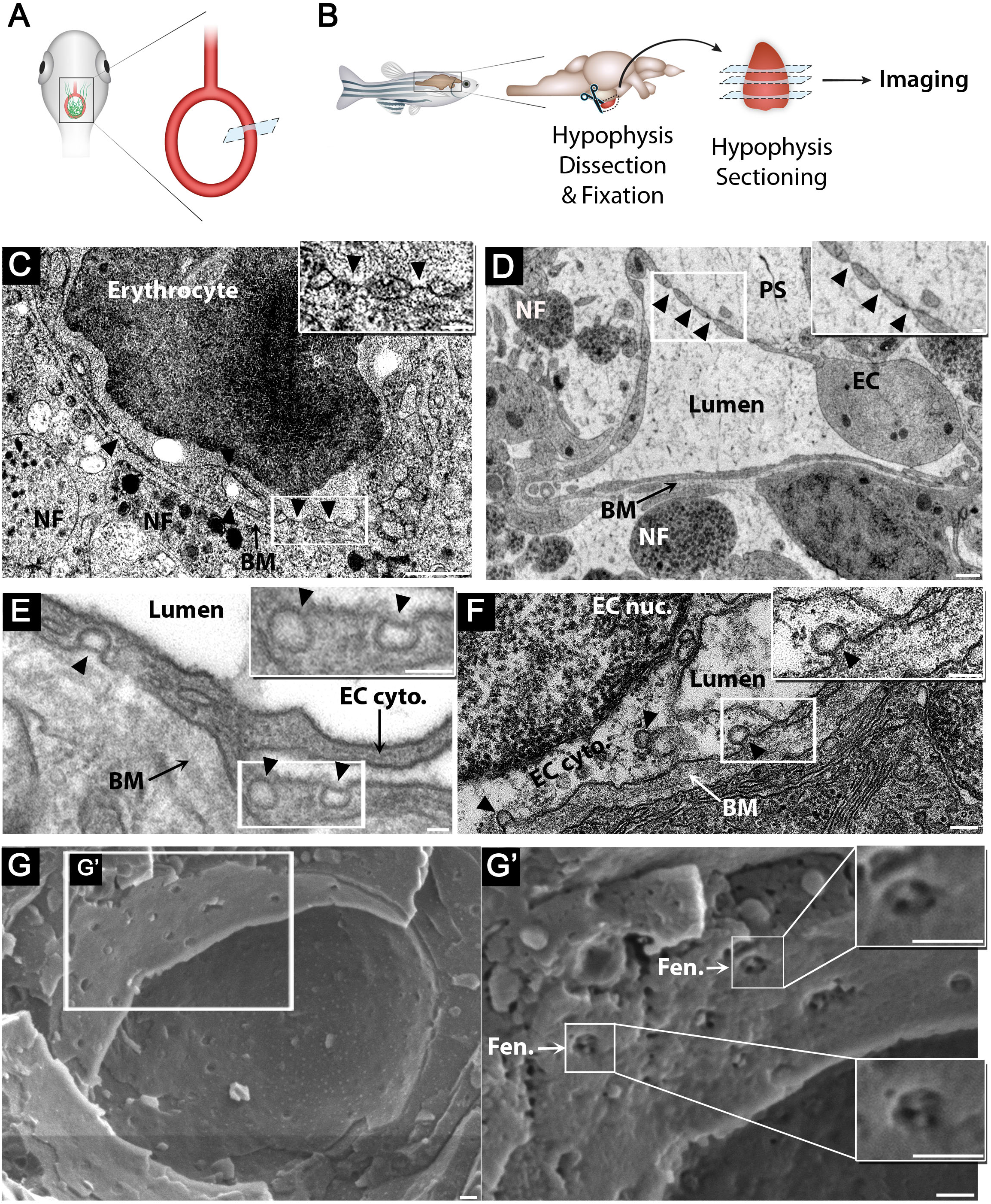
Zebrafish hypophyseal endothelia contain diaphragmed fenestrae and caveolae. (**A**) Scheme describing TEM imaging of the hypophyseal capillary in zebrafish larvae. The hypophyseal *oxtl*:EGFP+ cells was used as an anatomical landmark to localize the position of larval hypophysis prior to tissue preparation for TEM imaging. (**B**) Scheme describing TEM imaging of adult zebrafish hypophysis. Multiple ultrathin sections (60-80 nm) of the dissected hypophysis from adult zebrafish were submitted to TEM imaging. (**C,D**) TEM imaging of cross-section of larval (5 dpf) (**C**) and adult (**D**) hypophyseal fenestrated capillary contacting hypothalamic axonal nerve fibers (NF). Scale bars: 500 nm (C), 200 nm (D). Insets: higher magnification of diaphragmed fenestrae organized in a sieve-like manner. Fenestrae are denoted by arrowheads. Scale bars: 100 nm. (**E,F**) TEM imaging of hypophyseal endothelia containing abluminal, luminal and internalized caveolae. Caveolae are denoted by arrowheads. Scale bars: 100 nm. Insets: higher magnification of internalized and surface caveolae with stomatal diaphragm. Scale bars: 100 nm. (**G**) Cryo-SEM imaging of fenestrated endothelia in adult zebrafish hypophysis. (**G’**) Higher magnification of the hypophyseal fenestrated endothelial cell, demonstrating surface view of fenestral diaphragm dissected into several openings by fibrils that converge at the center of the pore (insets). Fen., fenestrae, NF, nerve fiber, PS, perivascular space; BM, basement membrane, EC, endothelial cell., EC nuc., endothelial cell nucleus, EC cyto., endothelial cell cytoplasm.

In addition to fenestrae, zebrafish hypophysis contained classical neurovascular interfaces including axonal swellings enriched with large dense core vesicles, which were detected near the endothelial basement membrane and perivascular space (Fig. 3C,D). Hypophyseal capillaries also contained caveolae. Open caveolar vesicles appeared on the luminal and abluminal sides of the capillary, whereas the ‘internalized’ cytoplasmic caveolar vesicles were localized inside the endothelial cell (Fig. 3E,F).

### Zebrafish hypophyseal capillaries express two *PLVAP* orthologs

We recently reported that 5 dpf zebrafish larvae express the *PLVAP* ortholog *plvapb* (a.k.a. *vsg1*) in the hypophyseal loop (Anbalagan et al., 2018). Our bioinformatic analysis revealed that zebrafish genome contains two gene orthologs, *plvapa*, previously named, *si:dkey-208k22.3* and *plvapb* (Fig. 4A). The predicted zebrafish Plvapb protein resembles the secondary structure of the mammalian protein, including three instead of two coiled-coil domains. The predicted Plvapa protein, on the other hand, displays a mild deviation, as it contains four coiled-coil domains and a longer C terminus (Fig. 4A, S1).

**Figure 4.**
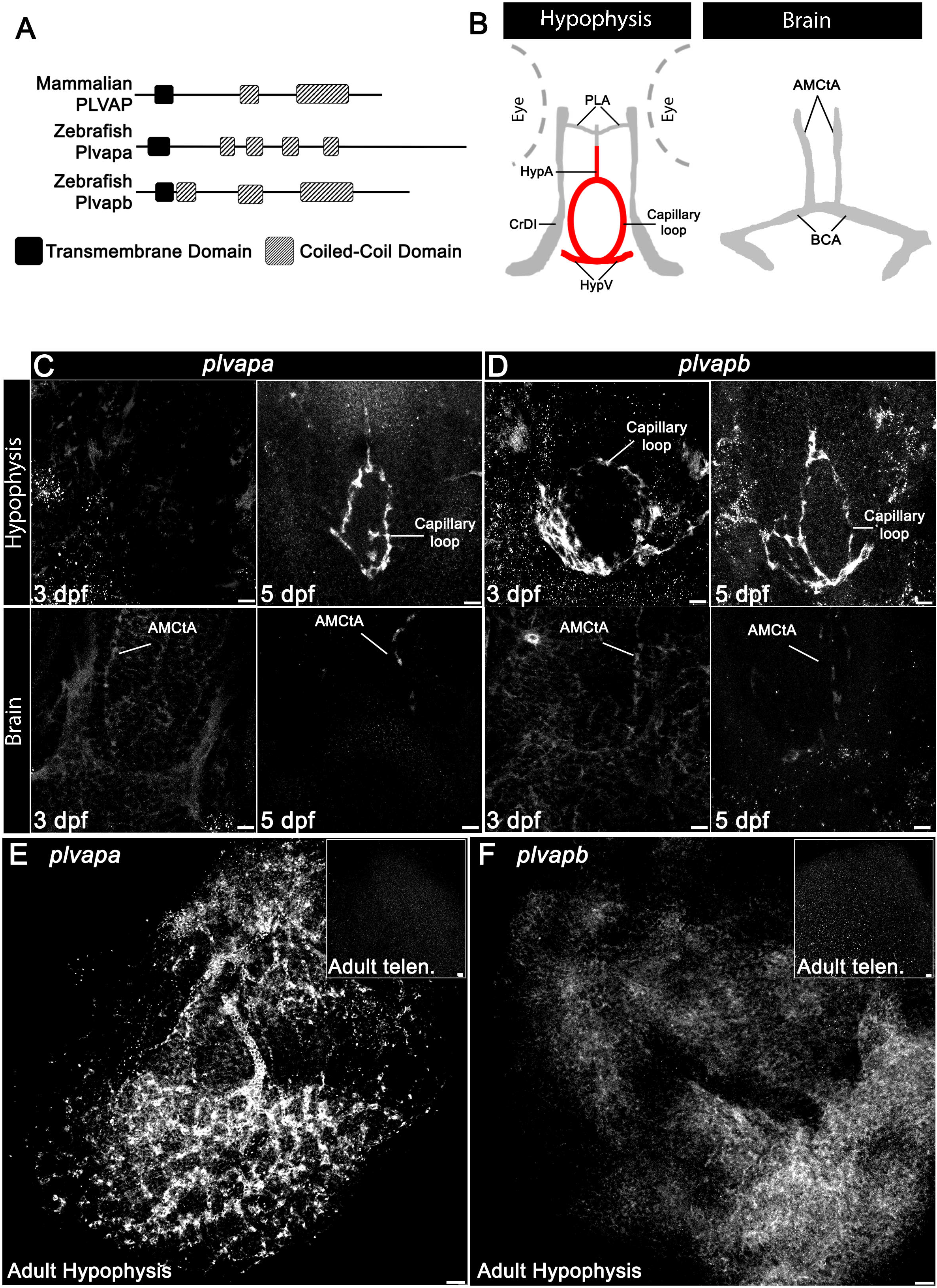
Developmental expression of Plvap orthologs in the hypophyseal vasculature. **(A)** Schematic representation of the predicted secondary structure of zebrafish Plvapa and Plvapb versus the mammalian PLVAP. The predicted zebrafish Plvapb protein resembles the secondary structure of the mammalian protein, but contains three instead of two coiled-coil domains. The predicted Plvapa protein displays a mild deviation, as it contains four coiled-coil domains and a longer C terminus. **(B)** Scheme describing zebrafish larval hypophyseal and brain vasculature. CrDI, Cranial division of the internal carotid artery, HypA, hypophyseal artery, AMCtA, Anterior (rostral) mesencephalic central artery, BCA, Basal communicating artery, PCS, Posterior (caudal) communicating segment. **(C-D)** Whole-mount fluorescent in situ hybridization (FISH) of zebrafish larvae at 3 or 5 dpf showing restricted *plvapa* (**C**) and *plvapb* (**D**) mRNA expression in the hypophyseal but not in the brain vasculature. AMCtA, Anterior (rostral) mesencephalic central artery. Scale bars: 5 μm. (**E-F**) Whole-mount FISH of adult zebrafish showing restricted *plvapa* (**E**) and *plvapb* (**F**) mRNA expression in the hypophyseal but not in the brain vasculature. Adult telen., adult telencephalon. Scale bars: 20 μm. See related Figures S1 and S2.

To explore the role of Plvap in hypophyseal endothelium, we examined the expression of *plvapa* and *plvapb* mRNAs by whole-mount fluorescent *in situ* hybridization (FISH) at 3 dpf, at 5 dpf, which coincides with the developmental acquisition of hypophyseal vessel permeability, and in adult fish. The expression of *plvapb* mRNA appeared in the hypophyseal vascular loop already at 3 dpf, while *plvapa* expression was only detected at 5 dpf (**Fig. C,D**). The expression of both genes persisted in adult hypophyseal capillary plexus (Fig. 4E,F). Whole-mount FISH on the background of transgenic Tg(*kdrl*:EGFP) zebrafish larvae (5 dfp) showed that the expression of both *plvapa* and *plvapb* mRNA is endothelial-specific (Fig. S2). In agreement with previous reports that PLVAP expression is down-regulated in brain endothelia upon BBB formation (Hallmann et al., 1995; Umans et al., 2017; van der Wijk et al., 2019), expression of *plvapa* and *plvapb* was not detected in larval and adult brain vasculature (Fig. 4C,D, bottom; 4E,F, insets).

### Mutation in *plvapb* leads to deficits in fenestral and stomatal diaphragms

To study the role of this gene in zebrafish we used a *plvapb* mutant, *plvapb^sa13080^*, which contains a point mutation leading to a premature stop codon, which results in a truncated Plvapb protein that has deficient functionally important extracellular coiled-coil domains (Fig. S3). Homozygous *plvapb^−/−^* zebrafish mutants were viable, allowing testing the role of Plvapb in fenestrae formation in the hypophyseal endothelia. Quantitative TEM analysis of hypophyseal sections from adult *plvapb^−/−^* fish showed a significant loss of both fenestral (Fig. 5A-C) and stomatal (Fig. 5D-F) diaphragms. However, some diaphragms were still evident in the fenestral openings, probably due to functional redundancy between *plvapa* and *plvapb* (Fig. 5B, black arrowheads). *plvapb^−/−^* fish displayed no differences in diaphragm thickness (Fig. 5G) of fenestrae and caveolae (Fig. 5H,I). Likewise, the mean diameter (Fig. 5G) of either fenestral or caveolar pores, with or without diaphragms, did not differ significantly between wild-type (WT) and mutant fish (Fig. 5J,K).

**Figure 5.**
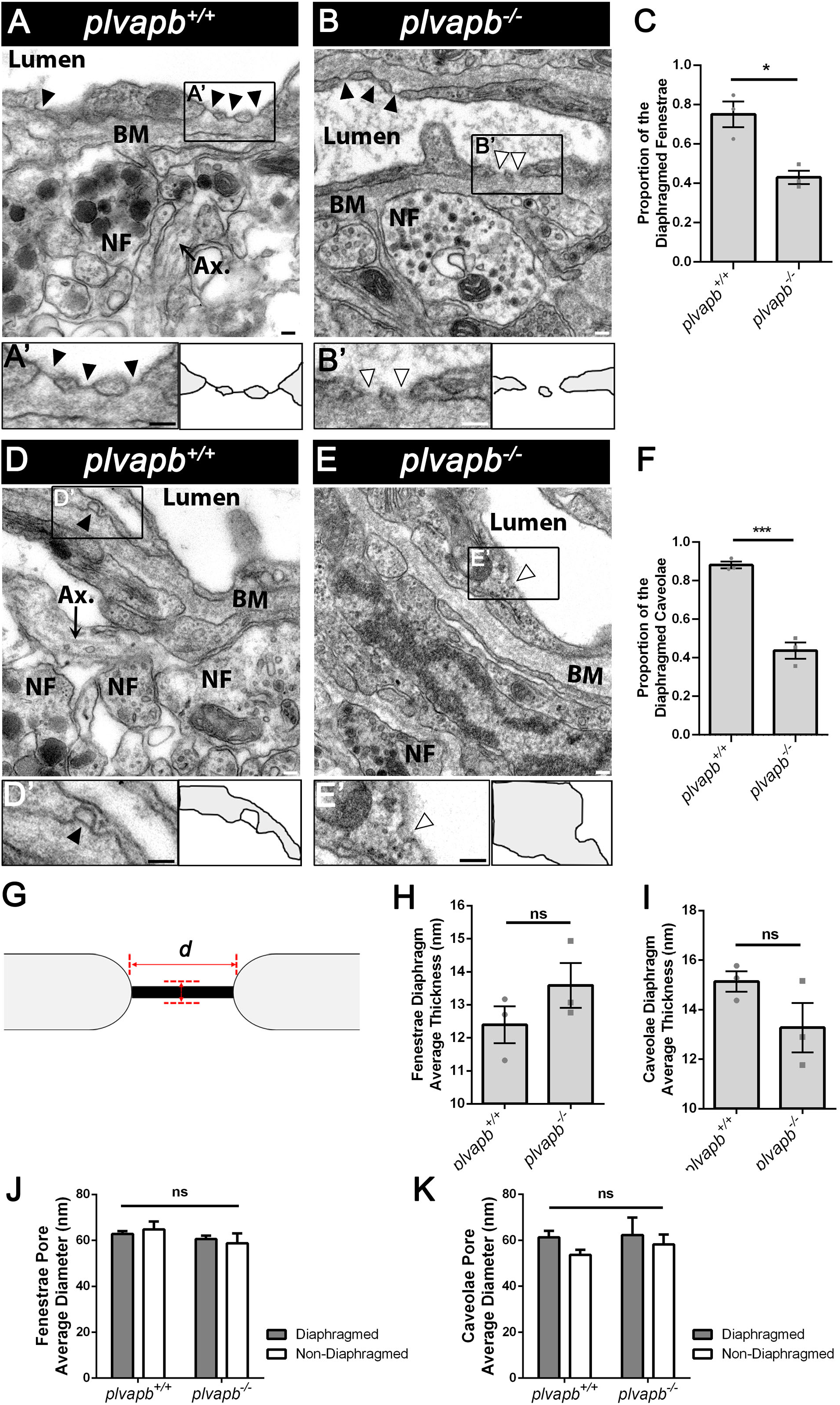
Ultrastructural analysis of fenestral and stomatal diaphragms in WT versus *plvapb^−/−^* zebrafish hypophyseal endothelium. (**A-C**) Ultrastructural analysis of fenestral diaphragms in adult zebrafish. The total number of diaphragmed and non-diaphragmed fenestrae was counted in each image; 10-45 images were obtained per specimen. Then, the proportion of complete diaphragms was calculated for each image, averaged for each specimen and then averaged for each genotype group; in total, ~60-150 fenestrae were analyzed per genotype. (**A**) TEM imaging of diaphragmed fenestrae (black arrowheads) in the *plvapb*^+/+^ adult zebrafish hypophysis. (A’) Higher magnification and graphical depiction of diaphragmed fenestrae. (**B**) TEM imaging of non-diaphragmed fenestrae (white arrowheads) in the *plvapb^−/−^* adult zebrafish hypophysis. Some diaphragmed fenestrae (black arrowheads) were still evident in the *plvapb^−/−^* mutants. (**B’**) Higher magnification and graphical depiction of non-diaphragmed fenestrae. (**C**) Graph showing the proportion of diaphragmed fenestrae in *plvapb*^+/+^ versus *plvapb^−/−^* mutant fish. The proportion of complete diaphragms was significantly decreased in the mutant fish (*p<0.05; Welch two-sample *t*-test, n=3 for each genotype). (**D-F**) Ultrastructural analysis of caveolar diaphragms in the adult zebrafish hypophyseal endothelia, quantified as in (**A-C**). 10-45 images were obtained per specimen, ~70-120 caveolae were analyzed per genotype group. (**D**) TEM imaging of diaphragmed caveolae (black arrowheads) in the *plvapb*^+/+^ adult zebrafish hypophysis. (**D’**) Higher magnification and graphical depiction of diaphragmed caveolae. (**E**) TEM imaging of non-diaphragmed caveolae (white arrowheads) in the *plvapb^−/−^* adult zebrafish hypophysis. (E’) Higher magnification and graphical depiction of non-diaphragmed caveolae (**F**) Graph showing the proportion of diaphragmed caveolae in *plvapb*^+/+^ versus *plvapb^−/−^* mutant fish. The proportion of complete stomatal diaphragms was significantly decreased in the mutant fish (***p<0.001; Welch two-sample *t*-test, n=3 for each genotype). (**G-K**) Quantitative analyses of fenestral and caveolar diaphragms ultrastructural parameters. (**G**) Scheme describing how the diaphragm thickness and pore diameter of fenestrae and caveolae were measured. (**H,I**) Graphs showing mean fenestral (**H**) and caveolar (**I**) diaphragm thickness. The thickness of each diaphragm was measured three times using a line drawing tool in ImageJ and averaged. The mean value of the measurements was calculated for each specimen and then averaged for each genotype group. In total, ~40-100 fenestral and stomatal diaphragms were analyzed in the *plvapb*^+/+^ specimens and ~35-66 fenestral and stomatal diaphragms were analyzed in the *plvapb^−/−^* specimens. No significant difference were found between *plvapb*^+/+^ and *plvapb^−/−^* fish (ns, not significant; Welch two-sample *t*-test, n=3 for each genotype) (**J,K**) Graphs showing mean fenestral (**J**) and caveolar (**K**) pore diameter. The diameter of each opening was measured three times using a line drawing tool in ImageJ and averaged. The mean value of the measurements was calculated for each specimen and then averaged for each genotype group. In total, ~40-100 fenestral and caveolar openings were analyzed in the *plvapb*^+/+^ specimens and ~60-150 in the *plvapb^−/−^* specimens. No significant difference in the diameter of either diaphragmed or non-diaphragmed fenestral and caveolar openings was found between the *plvapb*^+/+^ and *plvapb^−/−^* fish (ns = not significant; two-way ANOVA, n=3 for each genotype). Data are presented as mean ± SEM (C, F, H, I, J, K). BM, basement membrane, NF, nerve fiber, Ax., axonal element. Scale bars: 100 nm.

As mentioned, it has been suggested that Plvap is not required for fenestrae formation *per se* (Herrnberger et al., 2012; Stan et al., 2012). Therefore, focusing on hypophyseal vasculature, we examined whether developmental mutation in the *plvapb* ortholog affected fenestrae formation. Fenestrae were counted along the length of the endothelial wall in each TEM image of adult zebrafish and their density was calculated. The mean density of fenestrae was more than 3 fenestrae per 10 μm of endothelium in *plvapb^−/−^* mutant fish, whereas in control *plvapb*^+/+^ fish, the density was only about 1.5 fenestrae along the same endothelial wall length (Fig. 6A-B, E. Notably, whereas there were more abluminal than luminal caveolae in hypophyseal endothelia of both WT and mutant fish, the quantification of caveolar density along the endothelium revealed no significant difference between the genotypes (Fig. 6C-D,F).

**Figure 6.**
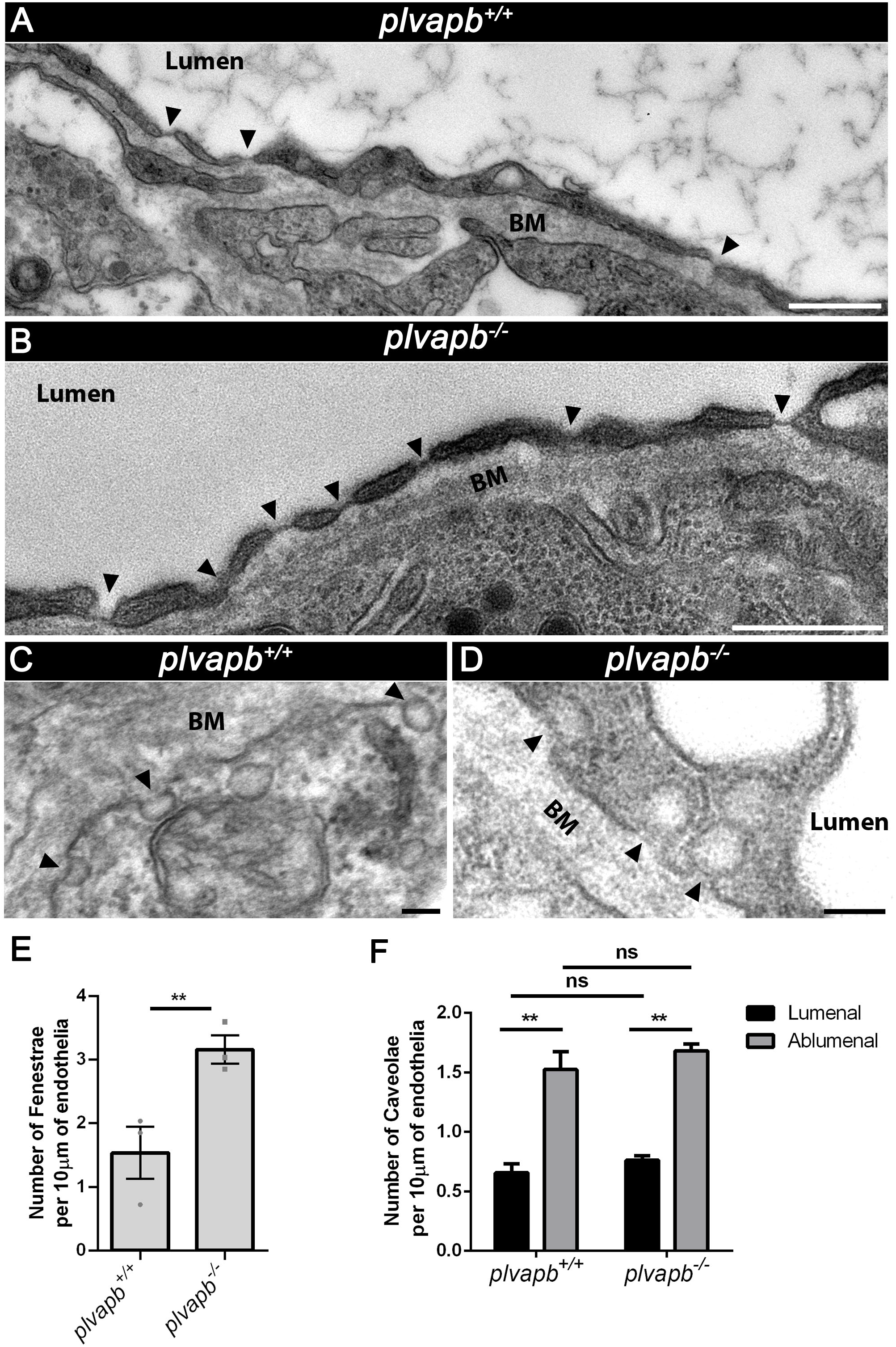
Mutation in *plvapb* leads to increased density of fenestrae but not caveolae. (**A,B**) Representative TEM images of endothelial cells of the *plvapb*^+/+^ (**A**) and *plvapb^−/−^* (**B**) adult zebrafish hypophyses, showing the increased number of fenestrae (arrowheads) along the endothelium in the mutant fish. Scale bars: 500 nm. (**C,D**) Representative TEM images of endothelial cells of the *plvapb*^+/+^ (**C**) and *plvapb^−/−^* (**D**) adult zebrafish hypophyses, showing multiple caveolae (arrowheads) along the endothelial wall. Scale bars: 100 nm. (**E**) Quantification of the linear density of fenestrae per length unit (10 μm) of endothelium. The number of fenestrae was counted and the endothelial cell length was measured in each image; 10-45 images were obtained per specimen. The linear density was calculated as number of fenestrae divided by endothelial wall length (in nm), averaged for each specimen and then averaged for each genotype group. The density was then multiplied by 10^4^ to present the result as density per 10 μm of endothelial length. A significant increase in fenestrae linear density was observed in the *plvapb^−/−^* specimens (**p<0.01; Students’ *t*-test, n=3 for each genotype). (**F**) Quantification of the linear density of abluminal and luminal caveolae per length unit (10 μm) of endothelial length. The number of luminal and abluminal caveolae was counted and the endothelial cell length was measured in each image; 10-45 images were obtained per specimen. The linear density was calculated, averaged and multiplied as described above. No significant difference was found between the *plvapb*^+/+^ and *plvapb^−/−^* fish (ns, not significant; two-way ANOVA, n=3 for each genotype). Data are presented as mean ± SEM (E, F).

Taken together, these results suggest that developmental loss of function of *plvapb* gene impedes the formation of fenestral and caveolar diaphragms in hypophyseal endothelia. Moreover, the increased fenestrae density following *plvapb* loss of function may imply the existence of a Plvap-dependent regulatory feedback loop.

### Plvap limits the rate of blood-borne protein transfer into the hypophysis

We hypothesized that loss of fenestral diaphragm would affect the barrier function of neurohypophyseal endothelia. To directly decipher the role of *plvapb* in the passage of blood-borne proteins into the hypophysis parenchyma, we employed the Tg(*l-fabp*:DBP-EGFP;*kdrl*:mCherry-caax) permeability reporter line (Fig. 1A). First, we examined the effect of loss of *plvapb* on hypophyseal vascular morphogenesis. Results showed no differences in the shape (roundness index) or total area of the hypophyseal capillary loop between *plvapb^−/−^* and control larvae (Fig. 7A-C). We next measured the accumulated fluorescence intensity of extravasated DBP-EGFP in the hypophyseal loop of fixed Tg(*l-fapb*:DBP-EGFP;*kdrl*:mCherry-caax) larvae, which were generated on the background of either *plvapb^−/−^* or its WT siblings (Fig. 7A). This analysis showed no significant differences between the WT and *plvapb^−/−^* mutant larvae (Fig. 7D), indicating that Plvapb does not affect the steady-state hypophyseal vascular permeability.

**Figure 7.**
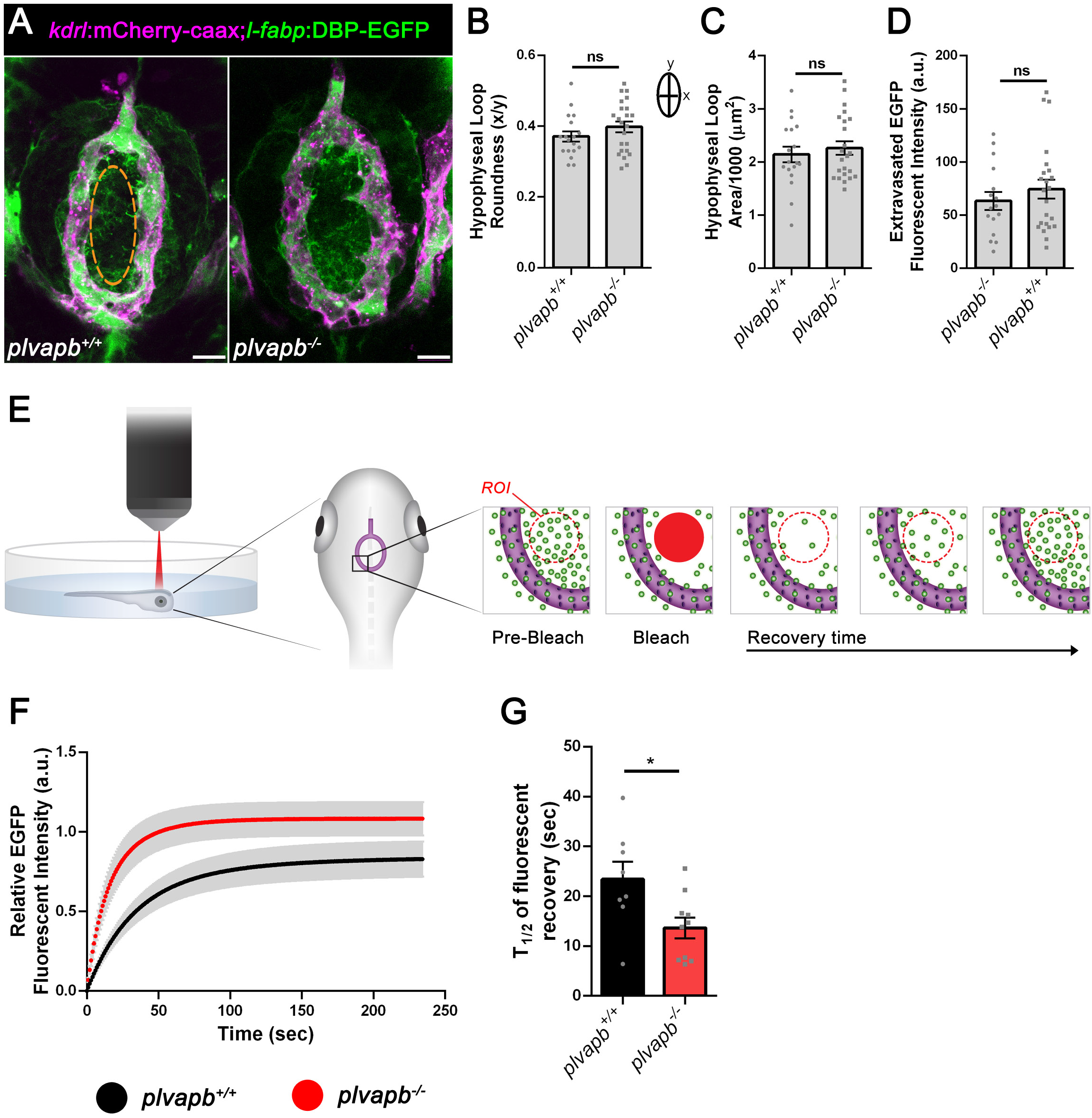
Plvapb limits the transfer rate of blood-borne proteins into the hypophysis parenchyma. (**A**) Confocal maximal intensity images showing the morphology of hypophyseal capillary loop in *plvapb*^+/+^ and *plvapb^−/−^* double transgenic Tg(*l-fabp*:DBP-EGFP;*kdrl*:mCherry-caax) zebrafish larvae (5 dpf). Scale bars: 10 μm. (**B,C**) Morphological analysis of hypophyseal capillary loop parameters, including roundness index (width (x)/ length (y)) (**B**) and area (**C**). No significant differences were found between *plvapb*^+/+^ (n=18) and *plvapb^−/−^* (n=23) fish (ns, not significant; Student’s *t*-test). (**D**) Quantification of the accumulated fluorescence intensity of the extravasated DBP-EGFP in the area inside the hypophyseal capillary loop (dashed line) of fixed *plvapb*^+/+^ (n=16) and *plvapb^−/−^* (n=23) larvae. No significant difference was found between the fish (ns, not significant; Student’s *t*-test). (**E**) Schematic representation of quantitative fluorescent recovery after photobleaching (FRAP) analysis of DBP-EGFP vascular extravasation in live zebrafish larval hypophysis. (**F**) The intensity of extravasated DBP-EGFP signal was measured in real time in live *plvapb*^+/+^ (n=8) and *plvapb^−/−^* (n=10) transgenic Tg(*l-fabp*:DBP-EGFP;*kdrl*:mCherry-caax) larvae (4 dpf) before and after bleaching. The time course of normalized EGFP fluorescent signal recovery after bleaching shows faster recovery in the *plvapb^−/−^* mutant. (**G**) Analysis of DBP-EGFP signal recovery half-time (T1/2), calculated by using ImageJ FRAP Profiler tool, shows faster blood-borne protein extravasation in *plvapb^−/−^* mutant versus WT fish (*p<0.05; Student’s *t*-test). a.u., arbitrary units; ROI, region of interest. Data are presented as mean ± SEM (B, C, D, F, G).

Next, we undertook real-time live imaging approach to measure the kinetics of DBP-EGFP entry into the hypophyseal parenchyma by performing quantitative fluorescent recovery after photobleaching (FRAP) analysis in live embryos. Thus, live transgenic Tg(*l-fapb*:DBP-EGFP;*kdrl*:mCherry-caax) larvae were subjected to FRAP analysis using a two-photon microscope. Specifically, a distinct region of interest (ROI), external to hypophyseal blood capillary, was photobleached and DBP-EGFP extravasation was monitored in real time (Fig. 7E). The intensity of EGFP signal was measured before and after the bleaching and the normalized fluorescent intensity was plotted as a function of time (Fig. 7F). The half-time of fluorescent signal recovery (T1/2), which was calculated using ImageJ FRAP Profiler tool (Fig. 7G), reflects the diffusion rate of the DBP-EGFP protein within the hypophysis parenchyma and, consequently, portrays the extravasation rate from the adjacent fenestrated blood capillary. Our FRAP analysis revealed that the recovery time of DBP-EGFP signal (T_1/2_) was less than half in the *plvapb^−/−^* mutants than in their WT siblings (13.63 ± 2.08 sec, n=10 versus 29.91 ± 7.19 sec, n=8, respectively), indicating a faster rate of DBP-EGFP extravasation in the absence of Plvapb (Fig. 7F,G).

Taken together, our results provide the first direct proof that PLVAP, a structural protein of the fenestral and stomatal diaphragms, regulates the rate of blood-borne protein passage through the fenestrated endothelial wall into the hypophyseal parenchyma.

## Discussion

Fenestrated endothelia containing PLVAP-positive diaphragm are found in various organs, which communicate with the blood circulation by rapid exchange of solutes, hormones and blood-borne proteins. Classical examples of such tissues are endocrine glands, intestinal villi, kidney glomeruli and circumventricular organs (CVO) in the brain (Aird, 2007; Miyata, 2015, 2017; Stan, 2007). Here, we investigated the role of PLVAP in regulating the passage of plasma proteins through the fenestrated endothelia of the hypophysis.

The deep brain location of the hypophysis and the fact that mammalian embryos develop *in utero* make it difficult to determine the developmental stage at which hypophyseal vascular permeability is established. Consequently, we lack information about the developmental acquisition of hypophyseal permeability. We therefore employed the genetically labelled permeability biosensor, DBP-EGFP, to visualize vascular permeability in the developing hypophysis of zebrafish. We found that a permeable hypophyseal capillary is established at a very early developmental stage and that this process is inversely correlated with the formation of BBB in CNS vasculature in the developing larvae. Notably, the hypophysis is regarded as one of several CVOs, which are permeable vascular interfaces devoid of BBB that surround the midline brain ventricles. These structures facilitate the molecular crosstalk between the brain and the peripheral circulation to regulate endocrine and metabolic function, thereby controlling homeostasis (Ganong, 2000; Miyata, 2017).

Previous studies have shown that the expression of Plvap becomes restricted to CVO vasculature upon formation of the BBB in other CNS blood vessels (Hallmann et al., 1995; Umans et al., 2017; van der Wijk et al., 2019). We show here that both *plvapa* and *plvapb*, the zebrafish orthologs of the mammalian *PLVAP*, are expressed by the embryonic hypophyseal endothelia and their confined pituitary expression coincides with the establishment of a sharp boundary between the permeable hypophyseal capillaries and the non-permeable brain vessels. What regulates the restricted *plvap* expression in the hypophysis and other CVOs is still unclear. We have recently shown that Vegf and TGFβ signaling are required to maintain vascular permeability as well as *plvap* expression in the developing hypophysis (Anbalagan et al., 2018). Inhibition of both TGFβ and VEGF signaling pathways, but not of either of them alone, causes regression of choroid plexus vascular fenestrations (Maharaj et al., 2008). Moreover, VEGF plays a major role in induction of endothelial fenestrations and in maintaining their integrity in mature tissues (Hamilton et al., 2019; Kamba et al., 2006; Stan, 2007). However, it is currently unclear why comparable levels of VEGF and TGFβ induce fenestrae formation only in some vasculatures but not in others, which are often adjacent. Interestingly, we found a temporal separation between *plvapa* and *plvapb* expression; whereas *plvapb* is expressed by hypophyseal endothelia already at 3 dpf, during the angiogenic development of the primary capillary loop, the expression of *plvapa* is visible only later in development. This suggests that *plvapb* might be active during the initial acquisition of hypophyseal vascular permeability.

It has been reported that Plvap is crucial for formation of fenestral and stomatal diaphragms; yet, it is not required for the formation of the fenestral pores themselves (Herrnberger et al., 2012; Ioannidou et al., 2006; Stan et al., 2012). We show here that *plvapb^−/−^* mutants exhibited significant loss of both fenestral and caveolar/stomatal diaphragms in the hypophysis. However, complete diaphragms were still found in the fenestrae and caveolae in these mutants. This phenotype is most likely due to functional redundancy between *plvapa* and *plvapb*, as both genes are expressed in the hypophyseal vasculature in both larvae and adult fish.

Interestingly, we found that *plvapb*-deficient fish display increased density of fenestrae but not of caveolae in the hypophyseal endothelia. This phenotype was not found in the mouse *Plvap* knock-out studies. Thus, the increased density of fenestrae in *plvapb^−/−^* mutant may imply the existence of a Plvap-dependent feedback mechanism, in which lower levels of Plvap protein activate an endothelial signaling cascade that drives fenestrae formation.

Several studies on *PLVAP* knock-out mice have shown that the blood composition of these animals is severely disrupted, leading to hypoproteinemia, hypoalbuminemia and hypertriglyceridemia as a result of severe non-inflammatory protein-losing enteropathy (Herrnberger et al., 2012; Stan et al., 2012). These results were recently supported by independent case studies of human infants with homozygous nonsense mutation in *PLVAP* gene. The patients (<0.5 years old) suffered from severe protein-losing enteropathy caused by the absence of fenestral and stomatal diaphragms and eventually died (Broekaert et al., 2018; Elkadri et al., 2015). These studies indicate that PLVAP protein plays a crucial role in maintaining barrier function of fenestrated endothelium, preventing uncontrolled leakage of plasma proteins from the blood vessels (Guo et al., 2016). However, thus far there was no direct demonstration that Plvap restricts the passage of substances through the endothelial wall. By directly measuring the *in vivo* kinetics of plasma protein extravasation from the hypophyseal capillaries in *plvapb^−/−^* mutant versus WT larvae, we now show that Plvap limits the rate of passage of DBP-EGPF through the fenestrated endothelial wall. The deficiencies in both fenestral and caveolar diaphragms and the increased density of fenestrae in the hypophyseal endothelial cells that we observed may contribute to the faster rate of plasma protein extravasation.

To conclude, we provide here the first direct proof that Plvap, a structural protein of the fenestral diaphragm, regulates the entry of plasma proteins into the hypophysis.

## Authors’ contributions

L.G., and G.L. planned the project. L.G. performed confocal imaging, FISH and immunofluorescent staining, established, performed and analyzed vascular permeability assay and FRAP, performed quantification and data analysis. J.B. generated the transgenic Tg(*oxtl*:EGFP) zebrafish line and contributed to the initial conceptualization of the project. L.G. and E.S. performed the TEM imaging. L.G. and D.G. performed the cryo-SEM imaging. B.A.A provided the transgenic Tg(*l-fabp*:DBP-EGFP) zebrafish line. G.L. and L.G. wrote the manuscript. All authors reviewed the manuscript’s text.

## Acknowledgments

We thank Roy Hofi for animal care; Vyacheslav Kalchenko for helping with the FRAP experiments; Ron Rotkopf for statistical analysis; Nitzan Konstantin for English editing; Tal Bigdary and Noa David for graphical schemes. The *plvapb^sa13080^* mutant was generated and provided by the Sanger Institute Zebrafish Mutation Project. B.A.A. is supported by the National Institute of Health (EY026181, EY027083, P30EY025585) and a Research to Prevent Blindness Challenge Grant. G.L. is supported by the Israel Science Foundation (#1511/16); US-Israel Bi-National Science Foundation (#2017325); Minerva-Weizmann program, Adelis Metabolic Research Fund and Yeda-Sela Center for Basic Research (in the frame of the Weizmann Institute). G.L. is an incumbent of the Elias Sourasky Professorial Chair.

## Materials and Methods

### Animal models

Zebrafish were raised and bred according to standard protocols. Embryos were raised at 28.5°C in 30% Danieau’s medium (0.17 mM NaCl, 0.21 mM KCl, 0.12 mM MgSO_4_, 0.18 mM Ca(NO_3_)_2_, 0.15 mM HEPES, pH 7.4) supplemented with 0.01 mg/L methylene blue. Prior to all labeling processes (detailed below), medium was supplemented with 8 mM PTU (1-phenyl-2-thiourea; Sigma-Aldrich, St. Louis, MO) to avoid formation of melanin pigments. The PTU was added at the beginning of gastrulation at ~5 hours post fertilization (hpf), and no later than 24 hpf and was kept in the medium until embryo fixation or live imaging.

### DNA extraction and genotyping

DNA for genotyping was attained from clipped fins of adult fish, 24 hpf embryos or from fixed samples after staining. DNA was extracted by submersion of the tissue samples in 50 mM NaOH, incubation for 40 minutes at 95°C, cool down for 5 minutes at 4°C, followed by neutralization with 1M Tris-HCl (pH 7.5; 1/10 of lysate volume). The lysate containing the genomic DNA was diluted 1:10 in ddH_2_O and used for DNA amplification. The genomic region of interest was amplified by using taqDNA Polymerase Master Mix Red 2x reaction mix (Ampliqon, Odense, Denmark). Amplified DNA region was detected by 0.8-1% agarose gel electrophoresis, then extracted and purified by using the NucleoSpin^®^ Gel and PCR Clean-up (Macherey-Nagel, Düren, Germany). Purified DNA fragments were sequenced by the Biological Services Unit at the Weizmann Institute of Science. Results were analyzed by ApE-A software (version 2.0, by M. Wayne Davis, University of Utah). See Table S1 for oligonucleotide sequences.

### Whole-mount fluorescent *in situ* hybridization (FISH) and immunofluorescent staining

Whole-mount zebrafish larva RNA FISH was performed as described in (Wircer et al., 2017). For RNA probe synthesis, partial coding sequences of the *plvapa* and *plvapb* genes were amplified by PCR, using the T7 polymerase (Roche) and purified with PCR clean-up kit. The purified products served as a template for synthesis of digoxigenin-labeled antisense mRNA probes using the DIG RNA labelling mix (Roche). The probes were then purified with RNAeasy mini kit (Qiagen, Hilden, Germany) and diluted in prehybridization solution (50% formamide in ddH2O, 5xSSC, 50 μg/ml heparin, 500 μg/ml yeast tRNA and 0.1% Tween-20, pH ~6.0) to a working concentration of 200 ng / 250 μl. See Table S1 for oligonucleotide sequences.

For immunofluorescent staining, larvae and freshly dissected adult hypophyses were first fixed overnight in 4% PFA and then washed in PBS (3×10 min). The samples were then permeabilized with PBSTx (Triton X100 0.5%; 3×20 min) and blocked in 500 μL of blocking solution (PBS + 10% goat serum + 1% DMSO + 0.5% Triton X100) for 1 hour at room temperature (RT). The solution was then replaced with 200 μL of fresh blocking solution with primary anti-Cldn5 antibody at 1:50 concentration and incubated overnight at 4°C. Samples were washed with PBSTx (4×30 min) and treated with 200 μL of secondary antibodies in blocking solution at 1:200 concentration overnight at 4°C. Then, samples were washed with PBST (Tween 0.1%; 3×30 min) and transferred to 75% glycerol (25-50-75%). Prior to imaging, the jaws were removed and the larvae were mounted ventrally on the slide. For antibody details, see Key Resources Table.

### Sequence analysis of the zebrafish Plvap orthologs

The analysis of Plvapa and Plvapb proteins was performed by using Expasy Proteomics tools server (https://www.expasy.org/proteomics). Specifically, the programs InterPro (http://www.ebi.ac.uk/interpro/) (Mitchell et al., 2018) and Phobius (http://phobius.sbc.su.se/) (Käll et al., 2004) were used for detection of transmembrane domains and COILS (https://embnet.vital-it.ch/software/COILS_form.html) for coiled-coil domains.

### Image acquisition

For TEM imaging of hypophyseal fenestrated endothelia, whole hypophyses from adult zebrafish were dissected and immediately frozen in HPM010 high-pressure freezing machine (Bal-Tec, Liechtenstein). Samples were subsequently freeze-substituted in an AFS2 freeze substitution device (Leica Microsystems, Austria) in anhydrous acetone containing 2% glutaraldehyde and 0.2% tannic acid for 3 days at −90°C and then warmed up to −30°C over 24 hours. Samples were washed in anhydrous acetone, incubated for 1 hour at RT with 2% osmium tetroxide and 2% uranyl acetate dissolved in ethanol. The samples were then washed with anhydrous acetone and infiltrated for 5-7 days at RT in increasing concentration of Epon in acetone.

For TEM imaging of hypophyseal fenestrated endothelia during early development, the transgenic Tg(*oxtl*:EGFP) zebrafish larvae (5 dpf) were first anesthetized with tricaine and small incision was made on the dorsal part of the scalp, followed by immediate fixation with fixative buffer (4% PFA, 0.2% glutaraldehyde, 0.1 M cacodylate and 5 mM CaCl_2_) overnight. The larvae were then embedded in 3.4% Noble Agar (DIFCO) and sectioned using vibratome (OTS-4000, Electron Microscopy Sciences, Hatfield, PA). EGFP-positive slices (~200 μm thick), which embrace the hypophysis were identified by fluorescent microscopy and were selected for further processing for TEM.

For quantitative ultrastructural analyses of fenestrae and caveolae TEM imaging of sectioned hypophyses from adult fish was performed. The whole hypophysis was dissected and immediately fixed in fixative buffer (4% PFA, 2% glutaraldehyde, 0.1 M cacodylate and 5 mM CaCl2) overnight. The adult whole hypophyses and larval EGFP-positive samples were then washed in 0.1 M cacodylate buffer and incubated in 1% osmium tetraoxide, 0.5% potassium dichromate, 0.5% potassium hexacyanoferrate in 0.1 M cacodylate for 1 hour. Samples were rinsed in 0.1 M cacodylate and then in ddH2O. Next, the samples were incubated in 2% uranyl acetate for 1 h covered with aluminum foil. Afterwards, samples were dehydrated in ethanol series and infiltrated for 5-7 days at RT in increasing concentration of Epon with ethanol.

The Epon-infiltrated samples, were polymerized in at 60°C for 48 hours. Ultrathin sections (60-80 nm) mounted a 200 mesh grids (Electron Microscopy Sciences, USA) supported with carbon-coated nitrocellulose film. The ultrathin sections double stained with 2% uranyl acetate in ddH2O and Reynolds lead citrate (Reynolds, 1963)and imaged by Tecnai T12 electron microscope operating at 120 kV, utilizing an ES500W Erlangshen CCD camera (Gatan, UK) or an Eagle 2K X 2K CCD camera (FEI).

For visualizing the fenestrae and caveolae, image montages of large areas that included either longitudinal or cross sections of endothelial cells were acquired with the Eagle CCD camera using the SerialEM program for TEM automated acquisition (Mastronarde, 2005). Alignment of montaged images was conducted using the “justblend” script included in the IMOD software package (Kremer et al., 1996).

For cryo-SEM, adult zebrafish hypophyses were dissected and fixed in 4% PFA and were then submitted to Cryo-SEM imaging as described in (Gur et al., 2018). In short, dissected pituitaries were cryo-immobilized in a high-pressure freezing device (HPM10; Bal-Tec). The frozen samples were transferred to a freeze-fracture device (BAF60; Bal-Tec), where they were freeze fractured and then coated with a 4-nm-thick layer of Pt/C. Samples were then observed by high-resolution SEM (Ultra 55, Zeiss) using secondary electrons and an in-lens detector, maintaining the frozen hydrated state by using a cryo-stage operating at a working temperature of −120°C.

Image processing was performed using ImageJ software.

### Fluorescent recovery after photobleaching (FRAP)

FRAP was performed in live Tg(*l-fabp*:DBP-EGFP;kdrl:) 4-days old WT and *plvapb^13080^* mutant larvae, using the LSM7 MP laser scanning microscope (Zeiss, Jena, Germany) with modified Achroplan X 40 0.8W, NA 1.0. The larvae were anesthetized with tricaine in Danieau’s solution without methylene blue. Larvae were then mounted in low melting agarose (1%) drop in a petri dish (60×15 mm) covered with Danieau’s solution. For bleaching, laser was set on 940 nm, 100% power for 20 iterations directed on multiple sequential small (~3 μm) regions of interest (ROIs) outside the hypophyseal capillary. For background signal, a region outside the pituitary was bleached as well. For reference signal, an additional region inside the pituitary capillary loop was measured, but not bleached. Consecutive time series images were taken before (n=10) and immediately after the photobleaching (290). Acquisition specifications were: Image dimension, 164×164 pixels (scaled: 37.96 μm x 37.96 μm); objective, W Plan-Apochromat 20×/1.0 DOC VIS-IR M27 75 mm; scan zoom, X-11.2, Y-11.2; pixel time, 1.23 μs; frame time, 38.96 ms. The raw datasets were first prefiltered with a Gaussian blur 3D (x=0, y=0, z=5), thereby averaging every five images (the first and last five images were excluded). The EGFP fluorescent signal data were then normalized to the background signal and a fitted curve was calculated per each specimen. The normalized EGFP fluorescent signal per time point was averaged for each genotype and plotted against relative time course. Average recovery half-time (T1/2) of the extravasated DBP-EGFP signal was calculated for WT and mutant groups. All the calculations were performed by ImageJ FRAP Profiler tool (http://worms.zoology.wisc.edu/research/4d/4d.html).

### Quantification and statistical analysis

Confocal images were analyzed using the open source ImageJ software. To quantify hypophyseal capillary loop morphology and permeability in fixed larvae, the Z-stacks encompassing the hypophyseal loop were summed up using ImageJ Z-project tool. To assess vascular permeability, the mean fluorescence intensity inside the loop area was measured (see Fig. 7A, dotted line). A region of interest of constant size outside the capillary loop was also measured for background correction. Specifically, the values of the background fluorescence were subtracted from the values of the internal capillary loop and averaged for each genotype group.

For the quantitative analyses of fenestrae and caveolae ultrastructure by TEM, the hypophyses from WT *plvapb*^+/+^ or *plvapb^−/−^* mutant fish were dissected and submitted to sample preparation and TEM imaging, as described above. The fenestrae and caveolae were identified by their unique fine structure and the pore diameter (50-80 nm for fenestrae; 50-100 nm for caveolae) according to previous ultrastructural studies on endothelial cells (Aird, 2007; Elfvin, 1965; Palade and Bruns, 1968; Rhodin, 1962; Stan, 2005; Stan, 2007).

For quantification, 10-45 images per specimen were analyzed by using ImageJ software. For ultrastructural analysis of fenestral diaphragms, the total number of diaphragmed and non-diaphragmed fenestrae was counted in each image. The proportion of complete diaphragms was calculated by dividing the number of diaphragmed fenestrae by the total number of fenestrae in the image. The proportion mean value was calculated for each specimen and the mean values of all specimens were averaged for each genotype group (n=3 for each genotype); in total, ~60-150 fenestrae were analyzed per genotype. For ultrastructural analysis of caveolae diaphragms, the total number of diaphragmed and non-diaphragmed caveolae was counted in each image. The proportion of complete diaphragms was calculated by dividing the number of diaphragmed caveolae by the total number of caveolae in the image. The proportion mean value was calculated for each specimen and the mean values of all specimens were averaged for each genotype group (n=3 for each genotype); in total, ~70-120 caveolae were analyzed per genotype.

For quantification of average fenestral and stomatal diaphragm thickness, the thickness of each diaphragm was measured three times using a line drawing tool in ImageJ and averaged. The mean value of the measurements was calculated for each specimen and the mean values of all specimens were averaged for each genotype group (n=3 for each genotype). In total, ~40-100 fenestral and stomatal diaphragms were analyzed in the *plvapb^+/+^* specimens and ~35-66 fenestral and stomatal diaphragms were analyzed in the *plvap^−/−^* specimens.

For quantification of average fenestral and caveolar pore diameter, the diameter of each pore was measured three times using a line drawing tool in ImageJ and averaged. The mean value of the measurements was calculated for each specimen and then averaged for each genotype group as described above (n=3 for each genotype). In total, ~40-100 fenestrae and caveolae were analyzed in the *plvapb^+/+^* fish and ~60-150 fenestrae and caveolae were analyzed in the *plvapb^−/−^* specimens.

For quantification of the linear density of fenestrae per length unit of endothelium, the number of fenestrae was counted and the endothelial cell length was measured within each image; 10-45 images were obtained per specimen. The density was calculated as the number of fenestrae divided by endothelial wall length (nm). The mean density was calculated for each specimen and then averaged for each genotype group (n=3 for each genotype). The density then was multiplied by 10^4^ to present the result as density per 10 μm of endothelial wall.

For quantification of the linear density of abluminal and luminal caveolae per length unit of endothelial wall. The number of luminal and abluminal caveolae was counted and the endothelial cell length was measured within each image, 10-45 images per each specimen. The density was calculated as number of caveolae divided by endothelial wall length (nm). The mean density was calculated within each specimen and then averaged with other specimens’ mean values within each genotype group (n=3 for each genotype). The density then was multiplied by 10^4^ to present the result as density per 10 μm of endothelial wall.

All data were tested for normality by Shapiro-Wilks test. Student’s *t*-test or Welch two-sample *t*-test was used for comparisons between two groups. Two-way ANOVA was used for comparison of pore diameters within diaphragmed and non-diaphragmed fenestrae and caveolae between the *plvapb^+/+^* and *plvapb^−/−^* fish, as well as for comparison of abluminal and luminal caveola density in *plvapb^+/+^* versus *plvapb^−/−^* fish. Statistical significance was determined as P<0.05. Data are presented as mean ± standard error of the mean (SEM).

---

**Figure S1.**
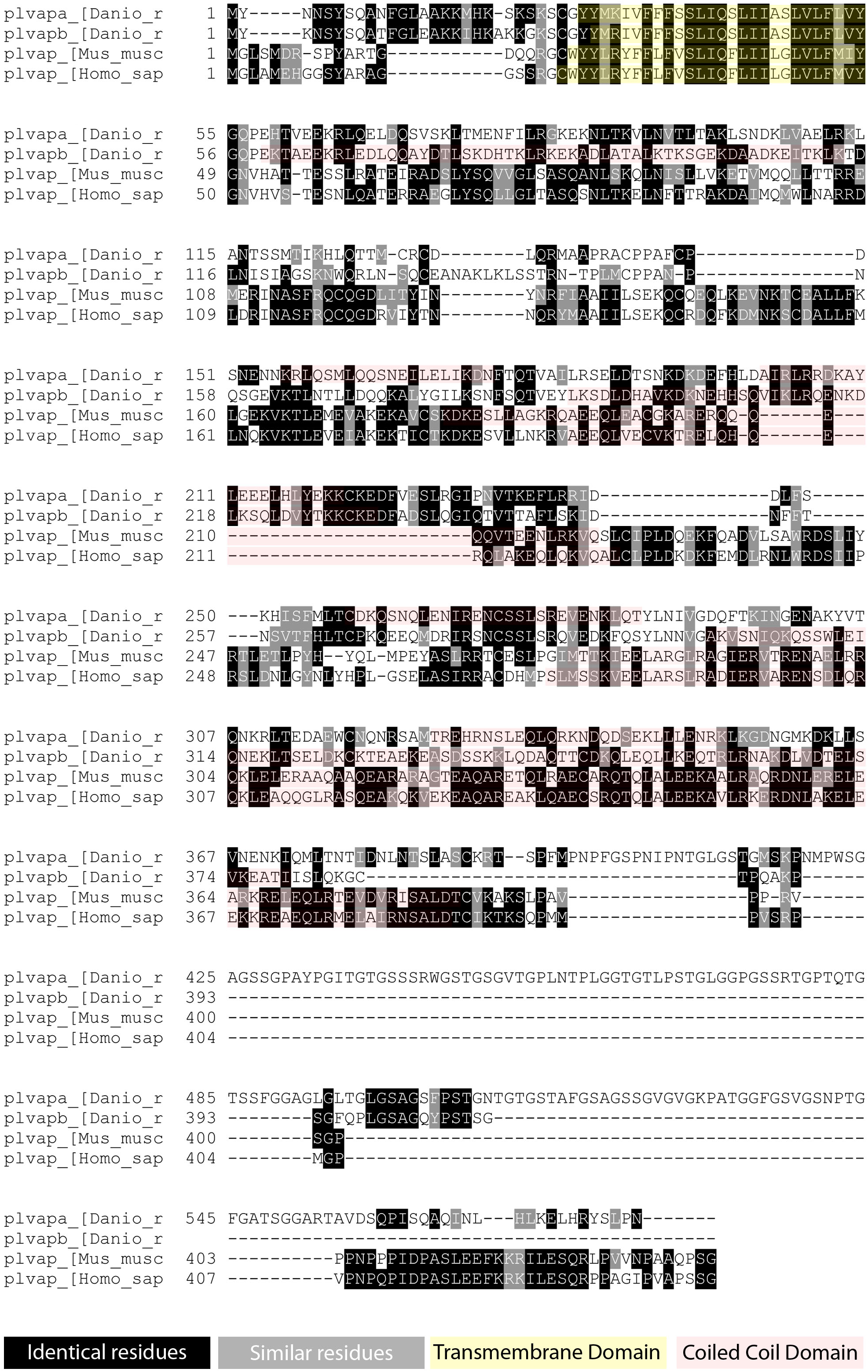
Multiple sequence alignment for Plvap protein from zebrafish versus mammalian species. Multiple sequence alignment of the deduced amino acid sequence of zebrafish proteins Plvapa (Plvapa_[Danio_r]] and Plvapb (Plvapb_[Danio_r]] compared to mice (plvap_[Mus_musc]) and human (plvap_[Homo_sap]) PLVAP proteins. Multiple sequence alignment was generated by Clustal 2.1, shaded according to the sequence identity between the species by BoxShade 3.21. Black shading indicates sequence absolute identity, gray shading indicates sequence similarity. The transmembrane domain is highlighted by yellow color, the coiled-coils are highlighted by red color.

**Figure S2.**
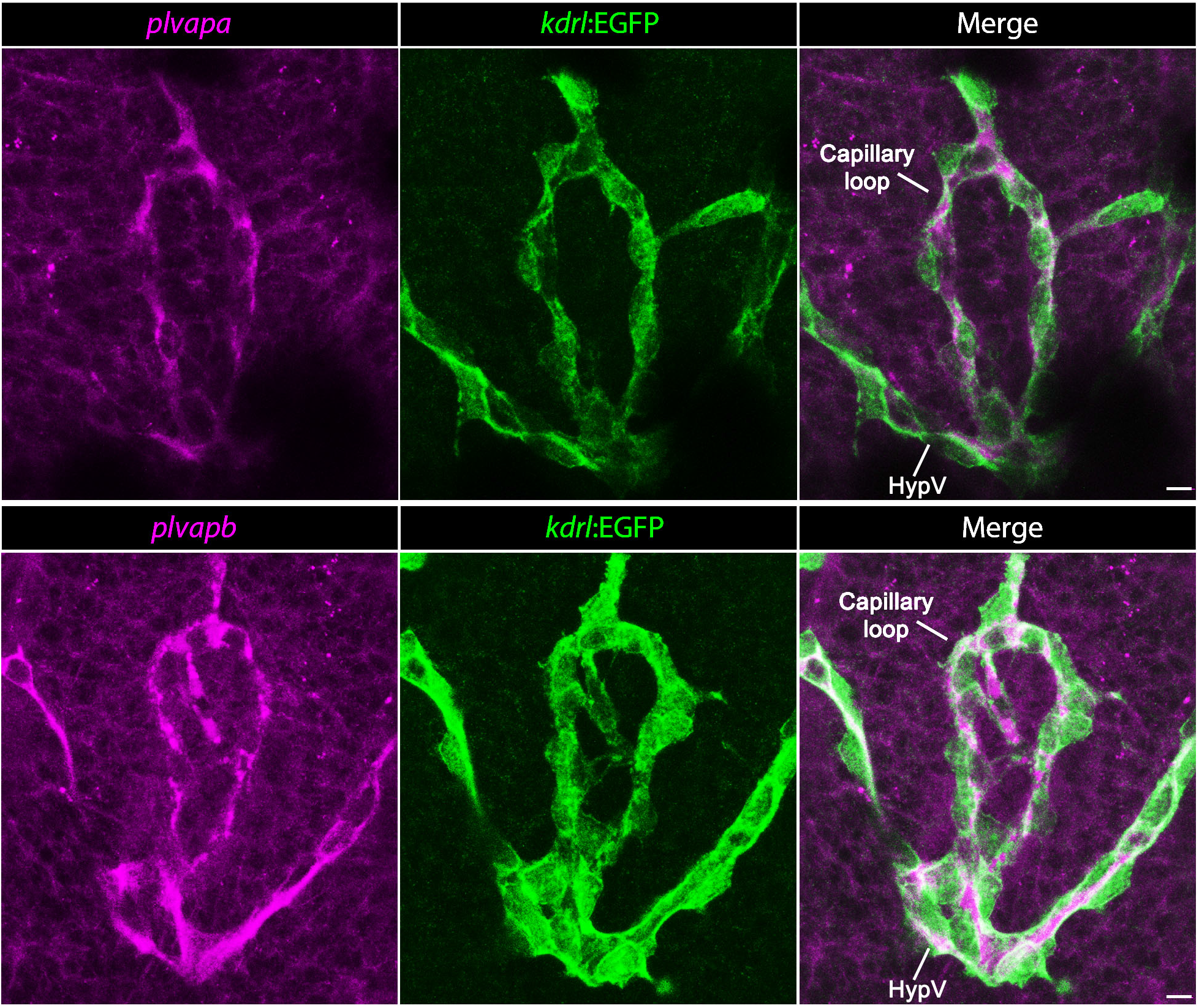
Zebrafish plvap orthologs are expressed in the hypophyseal vasculature. Whole-mount FISH of transgenic Tg(*kdrl*:EGFP) zebrafish larvae (5 dpf) showing *plvapa* and *plvapb* mRNA expression within the hypophyseal endothelial cells (green). HypV, hypophyseal vein. Scale bars: 5 μm.

**Figure S3.**
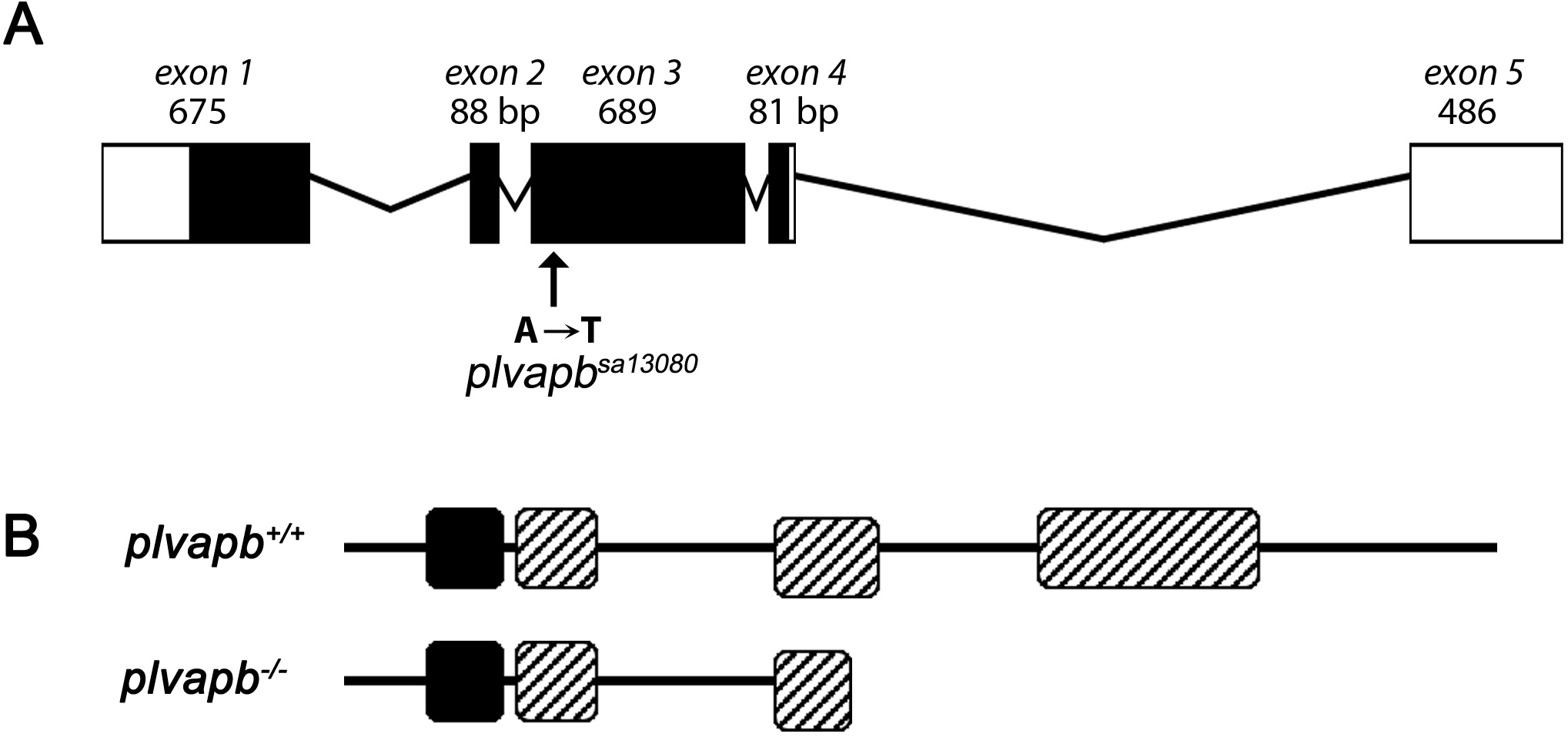
*Plvapb^−/−^* mutant description. (**A**) Schematic representation of the mutant *plvapb^sa13080^* allele including a nonsense point mutation (A→T) in the exon 3 of the zebrafish *plvapb* gene. (**B**) Schematic representation of the predicted secondary structure of *plvapb^−/−^* versus *plvapb*^+/+^ protein. In the mutant protein, the glutamine (a.a. 212) is replaced by premature stop codon, resulting in a shorter protein with only one functional coiled-coil domain.

**Table S1.**
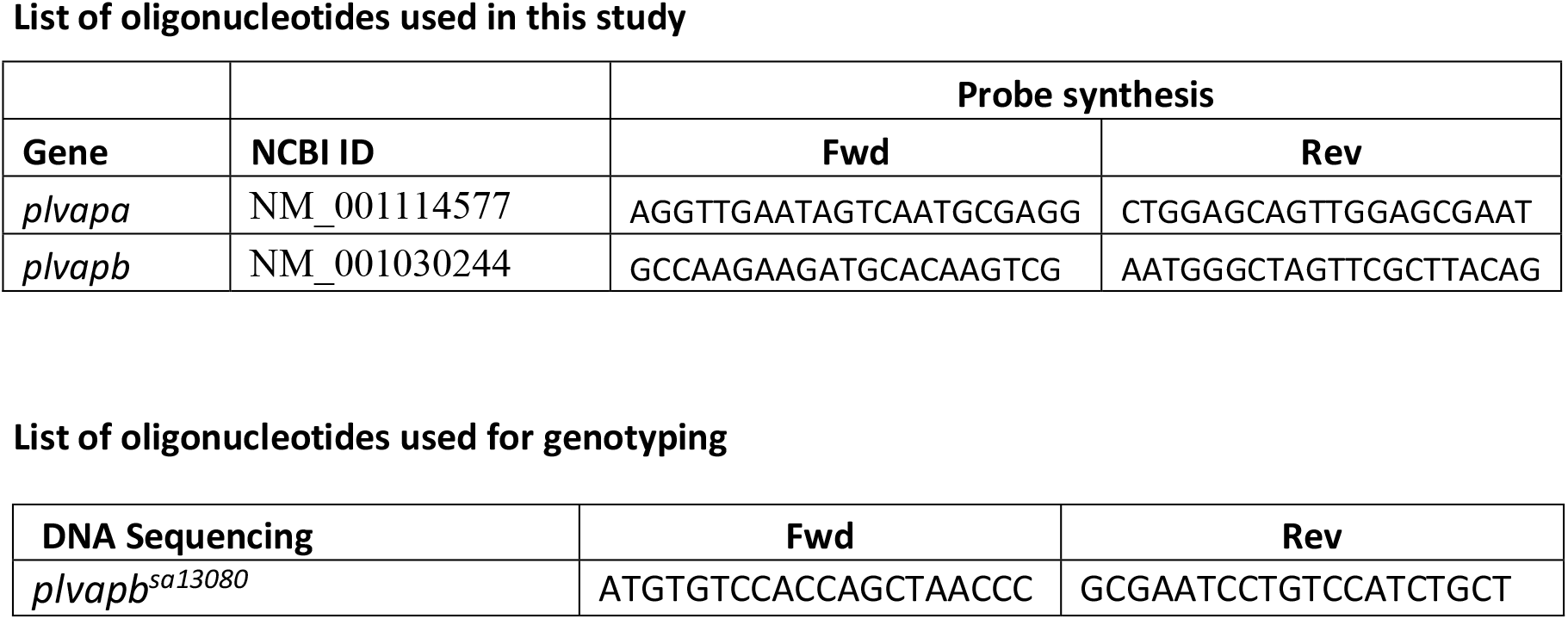

## KEY RESOURCES TABLE

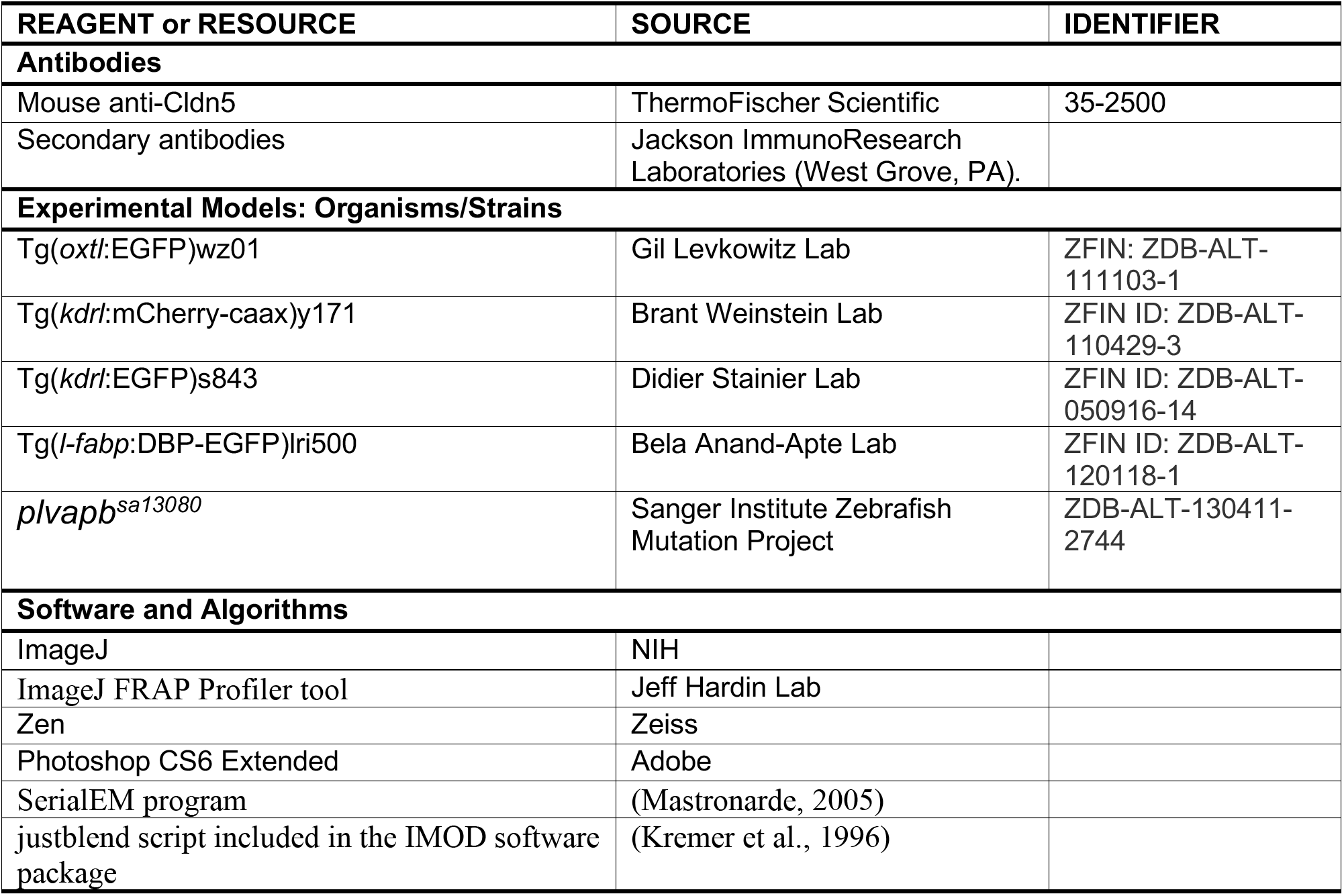

